# A genuinely hybrid, multiscale 3D cancer invasion and metastasis modelling framework

**DOI:** 10.1101/2024.01.12.575361

**Authors:** Dimitrios Katsaounis, Nicholas Harbour, Thomas Williams, Mark Chaplain, Nikolaos Sfakianakis

**Affiliations:** School of Mathematics and Statistics, University St Andrews, North Haugh, St Andrews, United Kingdom; School of Mathematical Sciences, University Nottingham, United Kingdom; School of Mathematics and Statistics, The University of Melbourne, Australia

**Keywords:** Cancer invasion, multiscale modelling, hybrid continuum-discrete, coupled partial and stochastic partial differential equations

## Abstract

We introduce in this paper substantial enhancements to a previously proposed hybrid multiscale cancer invasion modelling framework to better reflect the biological reality and dynamics of cancer. These model updates contribute to a more accurate representation of cancer dynamics, they provide deeper insights and enhance our predictive capabilities.

Key updates include the integration of porous medium-like diffusion for the evolution of Epithelial-like Cancer Cells and other essential cellular constituents of the system, more realistic modelling of Epithelial-Mesenchymal Transition and Mesenchymal-Epithelial Transition models with the inclusion of Transforming Growth Factor beta within the tumour microenvironment, and the introduction of Compound Poisson Process in the Stochastic Differential Equations that describe the migration behaviour of the Mesenchymal-like Cancer Cells. Another innovative feature of the model is its extension into a multi-organ metastatic framework. This framework connects various organs through a circulatory network, enabling the study of how cancer cells spread to secondary sites.

## 1. Introduction

Mathematical modelling has been a powerful tool in cancer research, aiding in the understanding of the intricate biological processes, in predicting cancer growth and metastasis, and in the treatment of the disease. However, the very dynamic nature of cancer, characterized by tumour heterogeneity, complex intercellular interactions, and evolutionary dynamics, presents challenges for traditional mathematical models. A particularly challenging aspect of cancer growth lies in the understanding of the intricate interplay between epithelial and mesenchymal phenotypes, which can greatly influence tumour progression and response to therapy.

In this context, we have previously proposed in [1, 2], a hybrid mathematical modelling framework that is capable of capturing the dynamic transitions between these phenotypes and shedding light on their underlying mechanisms. By combining deterministic and stochastic mathematical modelling frameworks, our hybrid approach is able to account for the heterogeneous nature of cancer cells, the complex macroscopic dynamics, and the stochasticity inherent in cellular migration. This integrated approach enables more comprehensive modelling of Epithelial-Mesenchymal Transition (EMT) and Mesenchymal-Epithelial Transition (MET) dynamics in cancer cells, cf. [3], offering insights into critical processes such as local tissue invasion, tumour island formation, and cancer metastasis. Our previously proposed model and work have provided us with valuable insights into the understanding of the relevant biological processes and the capabilities of the employed mathematics. It has also highlighted the direction of further model development.

In the current paper, we propose several model updates that better align with our accumulated understanding, incorporate new methodologies, and address the limitations of the original modelling framework, ultimately paving the way for even deeper insights, improved cancer predictions, and broader applicability in cancer modelling. The model extensions that we propose, incorporate among others porous medium-like diffusion for the time evolution of the Epithelial-like Cancer Cells (ECCs) and the various other living cell components of the system. Further model extensions account for more biologically realistic EMT and MET modelling by accounting for the role of Cancer-Associated Fibroblast (CAF) cells and Transforming Growth Factor beta (TGF-β) in the tumour microenvironment. In addition, the model features updated Stochastic Differential Equations (SDEs) for the migration of Mesenchymal-like Cancer Cells (MCCs). Most importantly though, we present here for the first time in our modelling approach, the progress we have made using our hybrid model on a multiple organ metastatic framework.

Mathematical modelling of cancer invasion goes back almost thirty years to the work of Gatenby and Gawlinski [4] and Perumpanani and colleagues [5]. These models were systems of nonlinear reaction-diffusion-taxis equations and studied travelling wave solutions modelling the invasive cancer cells cf. [6, 7]. Subsequent models considered hybrid discrete-continuum approaches [8], various discrete or individual-based approaches [9, 10], nonlocal modelling focussing on cell-cell and cell-matrix adhesion [11] and different multiscale approaches [12, 13, 14, 15, 16]. For a more comprehensive overview of the previous cancer modelling literature, the reader is referred to the recent review article of Sfakianakis and Chaplain [17].

The rest of the paper is structured as follows: in Section 2 we briefly describe the previous mathematical model, in Section 3 we present one after the other the proposed model extensions, Section 4 includes the complete updated model, and Section 5 accounts for a number of numerical simulations exhibiting aspects and properties of the overall updated model.

## 2. Previous hybrid model

The hybrid model that we present in this paper is an extension of a previously proposed hybrid model detailed in [1, 2]. Here, we provide a brief overview of the model.

The aforementioned hybrid model is comprised of two subsystems: the first incorporates the model components represented by their corresponding physical densities, including ECCs, ECM, and the matrix-degrading enzymes. The second subsystem focuses on the mesenchymal-like cancer cells describing them as individual/solitary migrating cells.

### 2.1 Density subsystem

We consider a Lipschitz set Ω ⊂ ℝ^2^ or ℝ^3^ and denote the (scalar) densities of the ECCs, ECM, and the matrix-degrading Metalloproteinases (MMPs) by *c*^*E*^(**x**, *t*), *v*(**x**, *t*), *m*(**x**, *t*) where **x** ∈ Ω respectively.

#### ECC density

The ECCs do not migrate actively. It is rather assumed, in this version of the model, that the colony of the ECCs disperses in the surrounding environment due to mechanical forces, e.g. internal pressure due to proliferation. This phenomenon is macroscopically captured through a (small in magnitude) diffusion term.

Furthermore, ECCs can be transformed into Mesenchymal-like Cancer Cells (MCCs) and vice versa by the EMT and MET, respectively. In this version of the model, it is assumed that EMT and MET are random and take place at a fixed rate. It is furthermore assumed that the ECCs proliferate in a logistic fashion, which accounts for the competition for free space among the ECCs the MCCs and the ECM macromolecules. These assumptions lead to the formulation of the following equation

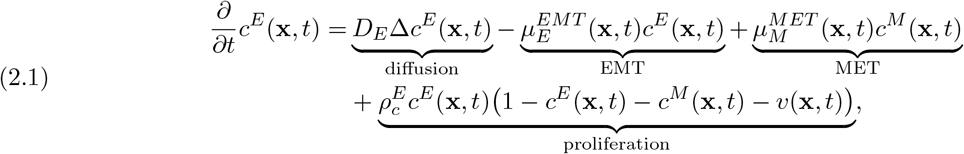

where 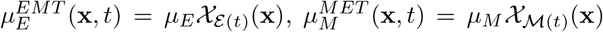, and with *D*_*E*_, *μ*_*E*_, *μ*_*M*_, 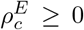 constants. Where 𝒳_*S*_(**x**) is the indicator function of the set *S* defined as:

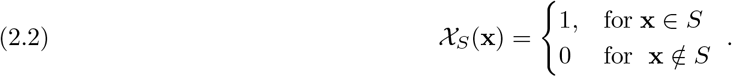

Here EMT occurs in a randomly chosen subset of the whole domain, ε(*t*) ⊂ Ω, with a fixed rate *μ*_*E*_. In a similar fashion, it is assumed that every (solitary) MCC might undergo MET independently from the others. The MCCs that undergo MET give rise to the subset ℳ(*t*) ⊂ Ω; MET is assumed to occur with a fixed rate *μ*_*M*_. Refer to Section 2.4 for a full discussion of the EMT and MET operators.

#### MMPs density

The MMPs are assumed to be produced by both types of cancer cells and to diffuse in the cancer microenvironment. It is also assumed that the MMPs decay at a constant rate. The time evolution of their density is given by the PDE:

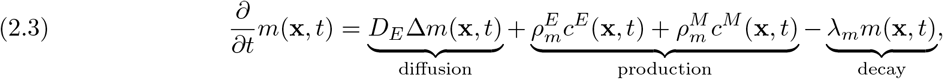

with constants 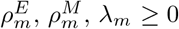.

#### ECM density

The ECM is represented by the density of collagen macro-molecules and it is modelled as a nonuniform and immovable component of the system that neither diffuses nor otherwise translocates. Additionally, it is assumed that the ECM is degraded by the action of the cancer-cell/MMP complex (both epithelial and mesenchymal). Finally, for the sake of model simplicity, it is assumed that the ECM is not reconstructed. The evolutionary equation of the ECM density is the following

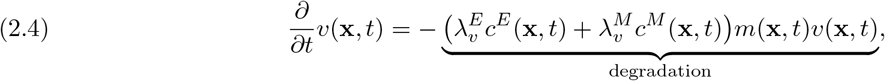

with constants 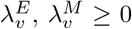. The assumption that the ECM is degraded by the cancer cell-MMP complex, rather than simply by the action of MMPs alone, is motivated by the fact that cancer cells require MT1-MMP, a specific type of membrane-bound MMPs, for invasion to take place [18].

### 2.2 Solitary cell subsystem

The MCCs are represented by a system of solitary cells indexed by *p* ∈ *P* = {1, 2, …, *N*}, *N* ∈ ℕ. Due to EMT and MET, the number *N* of solitary cells varies in time and, therefore, *N* = *N* (*t*). The MCCs are represented as point masses; **x**^*p*^(*t*) ∈ ℝ^3^ (or ℝ^2^) represents their position and *m*^*p*^(*t*) ≥ 0 their masses. Overall the set of solitary cells is described by

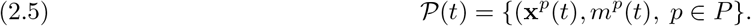

The migration of MCCs follows a biased random motion comprised of two independent processes: a directed migration part that represents the haptotaxis response of MCCs to gradients in the density of the ECM (drift term), and a stochastic component that represents the undirected kinesis of MCCs as they sense their environment, which is understood as Brownian motion (diffusion term).

The Brownian motion assumption is clearly a simplification which is justified by the random walk-like migration that the cells exhibit, see also [19]. Based on these assumptions, the migration of the solitary MCCs obeys the following SDE where 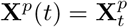 denotes the position of MCCs

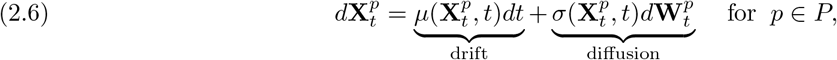

where 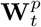 is a Wiener process with independent components. The drift and diffusion coefficients encode the modelling assumptions made about the nature of the directed and random components of the motion. We refer to [1] for more details regarding (2.6). With 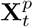 we denote the stochastic process describing the position of MCCs, satisfying SDEs of the form (2.6) and with **x**^*p*^ we denote the physical position of the solitary cancer cells represented by the set 𝒫(*t*) in (2.5).

#### Modelling reactions of MCCs

It is clear that (2.6) describes only the migration of the MCCs. Still, these cells participate in a number of reaction processes such as MET, EMT, production of MMPs, degradation of the ECM, and more.

These reaction terms are accounted for as follows: the MCCs undergo MET randomly, after which they are removed from the set *P* in (2.5) and transformed to a density, via the solitary-cell-to-density operator (introduced in Section 2.3), that is then added to the existing density of ECCs. Conversely, parts of the ECC density undergo EMT in a random fashion; this is initially transformed into MCC density, and subsequently, through a density-to-solitary-cell transition (introduced in Section 2.3), to solitary MCC cells. The newly formed solitary MCCs are then added to the existing set of solitary MCCs.

Besides EMT and MET, the full set of solitary MCCs is transformed into a density, denoted by *c*^*M*^ in (2.1), and subsequently employed in the evolution of the other components of the system that are described as densities. This is because the density of MCCs participates in the proliferation of ECCs (2.1), the production of MMPs (2.3) and the degradation of the ECM (2.4), which are all described via the density submodel.

### 2.3 Phase transitions between densities and solitary cells

The phase transition between the density description of the macroscopic ECCs and the solitary MCCs is conducted by the solitary-cell-to-density and density-to-solitary-cell operators. To proceed, it is assumed that the domain Ω is regular (e.g. a rectangle in two dimensions) and is sufficiently large to be split into equal computational partition cells *M*_*i*_

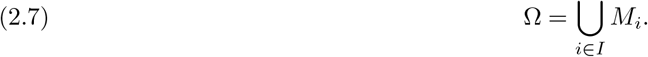

#### Density-to-solitary-cell transition

For a given density function *c*(**x**, *t*), the density-to-solitary-cell operator ℬ is given by

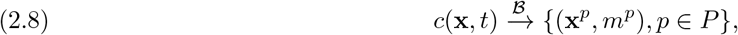

where at every partition cell *M*_*i*_ we assign cell with mass

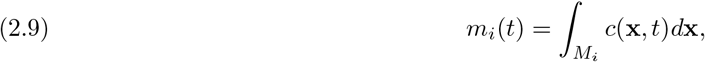

and position

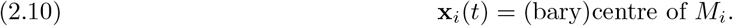

#### Solitary-cell-to-density

The solitary-cell-to-density operator ℱ is defined as

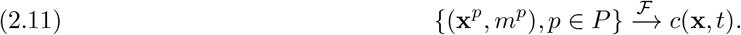

Every solitary cell is assigned to a rectangular domain *K*_*p*_ of size *K >* 0, centred at the centre of mass of the cells **x**^*p*^. It is assumed that the mass of the cell is evenly distributed over *K*_*p*_. As *K*_*p*_ overlaps with (possibly) several of the partition cells *M*_*i*_, we assign the corresponding portion of the cell mass given by

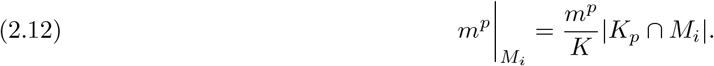

The mean value of *c*(**x**, *t*) over the partition cell *M*_*i*_ is denoted by *c*_*i*_(*t*). The density contribution of all solitary cells (*p* ∈ *P*) to the partition cell *M*_*i*_ as

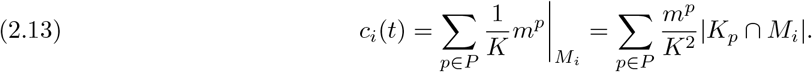

The density function across the full domain is then given by summing *c*_*i*_(*t*) over *i* ∈ *I*

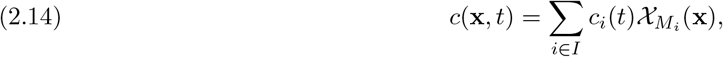

where 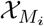 (**x**) is the indicator function defined by equation (2.2).

### 2.4. The EMT and MET operators

In this section, we present the mathematical description of the EMT and MET processes which, along with the density-to-solitary cell and solitary-cell-to-density operators, allow for coupling between the two cancer cell phenotypes.

#### EMT operator

It is assumed that a randomly chosen portion of the ECC density, 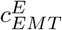, undergoes EMT to give rise to a density of MCCs

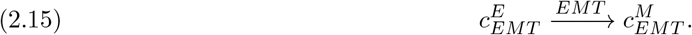

This newly created MCC density 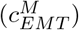 is then immediately transformed into solitary MCCs via the density-to-solitary-cell operator, described in Section 2.3

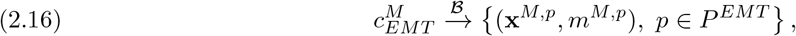

where **x**^*M,p*^, *m*^*M,p*^ are the position and mass of the newly created solitary cells and *P*^*EMT*^ is the corresponding set of indices. Following this, the collection of existing solitary MCCs is updated with the addition of the newly created solitary MCCs. This is given by the disjoint union

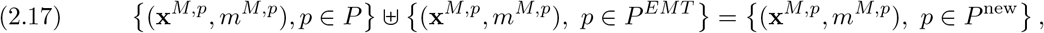

where *P* ^new^ is the re-enumeration of the set 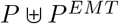. Overall, the EMT operator reads as (2.18)

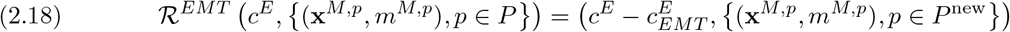

#### MET operator

It is assumed that each of the solitary MCCs undergoes MET randomly, to produce a set of solitary ECCs

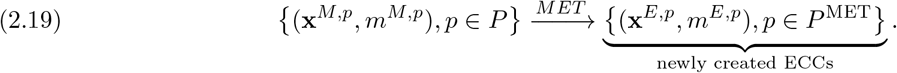

Subsequently, the solitary ECCs are transformed into a density via the solitary-cell-to-density operator

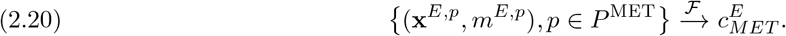

Overall, this can be written in operator form as

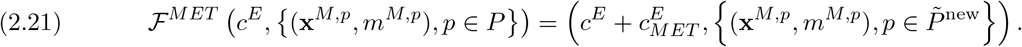

## 3. Extensions to the previous model

Having completed the presentation of the existing model, we move to the discussion of the proposed model updates. These updates encompass several key modifications including the incorporation of nonlinear diffusion—porous medium type—in the time evolution equations of the various cell types. Additionally, we address the role of CAF cells and of TGF-β in the tumour microenvironment. Furthermore, we refine the modelling of the EMT and MET operators and of the SDEs describing the migration of the MCCs. Lastly, we broaden the scope of our hybrid model by extending it to a multiple-organ metastasis framework, thereby facilitating a more comprehensive understanding of the progression and dissemination of cancer across the whole organism.

### 3.1 Nonlinear diffusion

The equation for the ECC evolution in the previous version of the model, (2.1), employed linear diffusion which (conditionally) leads to infinite propagation speed that the solutions exhibit. This implies that ECCs could spread instantaneously across the tissue, contradicting the actual biological nature of cancer cell migration, especially considering the timescale of cell migration and tumour growth. To address this discrepancy, our revised model introduces a simple nonlinear diffusion term, effectively resolving this issue.

In general terms, the diffusion functions *D*(*u*) in the nonlinear diffusion equation 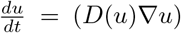 can be categorised into two main classes: degenerate and non-degenerate. Degenerate diffusion functions are characterised by their degeneracy at zero, i.e., *D*(0) = 0. The degeneracy ensures that, for any compactly supported initial condition, there is no diffusion at the interface where *u* = 0. As a result, the compactness of the initial conditions is maintained at all times. On the other hand, non-degenerate diffusion functions, such as linear diffusion, do not possess the property *D*(0) = 0. As a result, they do not create a sharp interface, and under certain conditions, can lead to infinite propagation speed. For a detailed discussion and comparison between degenerate and non-degenerate diffusion functions, we refer to [20].

This model extension, incorporating nonlinear degenerate diffusion functions, was first explored in [21], where various such functions were examined. Building on the remarks therein, we propose here the use of a Porous Medium type Equation (PME). The PME represents a natural extension of the classical heat/diffusion equation and is given in general form as

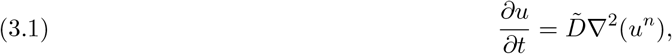

where *n >* 1. Assuming no-negative initial conditions, *u* remains non-negative for all times. Such would be the case in biological settings where *u* represents cell density, the PME (3.1) can be written as

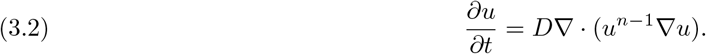

Clearly, the PME is degenerate as *D*(0) = 0 and admits solutions with a clear interface, compact support, and finite propagation speed. To demonstrate this key difference in behaviour we have plotted in Figure 1, typical solutions for both the Heat Equation and the PME. The solution in the Heat Equation spreads asymptotically in space, observing, hence, *infinite propagation speed*. In the contrary, the PME develops a distinct interface, with steep sides that propagate with finite speed. The PME, as its name suggests, was originally used to describe the flows through a porous medium, but has since been used in many other contexts [20]. In particular, porous-medium diffusion has been found to fit well with experimental data of biological cell migration [23, 24]. That is, it removes the problem of *infinite propagation speed* while introducing minimal additional complexity to the existing model.

**Fig. 1:**
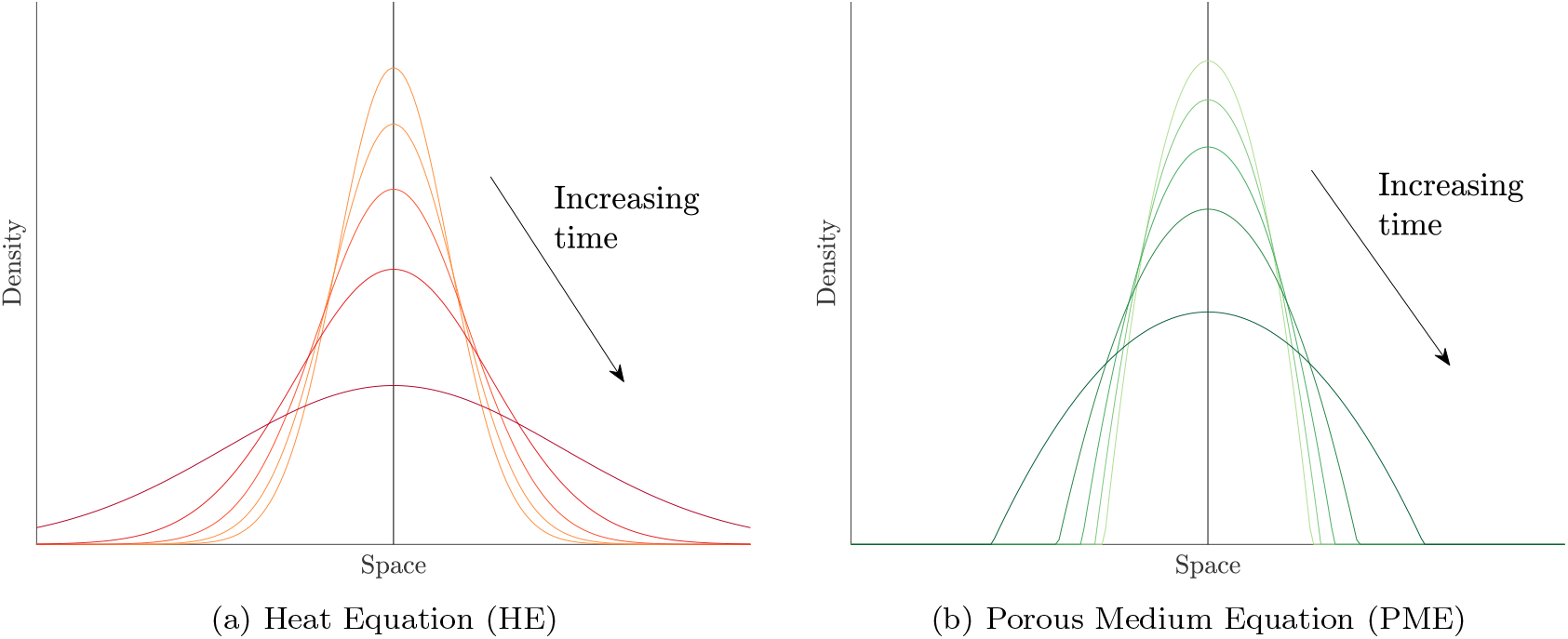
Numerical solutions of the (a) HE and (b) PME over time. A relative examination highlights the compactness of the solution’s support of the PME, justifying thereby the finite propagation observed by the PME solution. Figure source [21, 22].

**Fig. 2:**
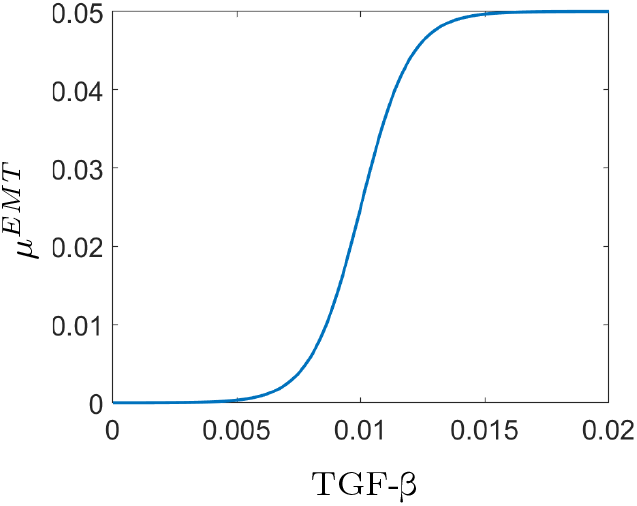
Graph of the EMT function *μ*^*EMT*^ (3.9) as a function of TGF-β for *L* = 0.05, *k* = 10^3^, and *b*_*T*_ = 0.01. Figure inspired from [30].

#### ECCs density with porous-medium diffusion

We introduce the nonlinear degenerate porous-medium type diffusion into the equation for ECC density. To this end, we consider the nonlinear porous-medium type diffusion function to be given by

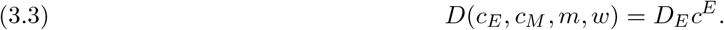

This is the simplest type of porous diffusion function and corresponds to *n* = 2 in (3.2). Using a diffusion function of this form, we implicitly assume that diffusion increases linearly with density. This is desirable as we have already stated that we assume ECCs primarily diffuse due to mechanical forces (e.g. pressure) caused by proliferation.

We retain all the original assumptions from (2.1). However, we replace linear diffusion with the porous-medium type diffusion function given in (3.3). Additionally, we assume that the ECCs compete with the CAFs which are included in the updated model and are introduced in Section 3.2. We have also included a haptotaxis term that accounts for the biased motion of ECCs, which is known to be in the direction of increasing ECM density [25, 26]. Therefore, the updated equation for ECC density is given by

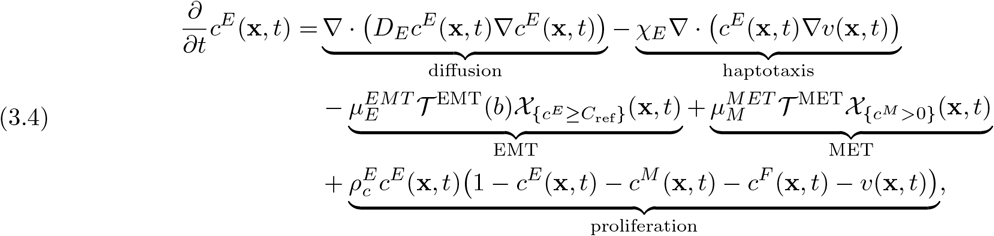

with constants 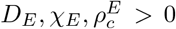. The EMT and MET operations are modelled in this current version as random processes through the variables 𝒯^EMT^ and 𝒯^EMT^. Moreover, in contrast to the original modelling in [1], EMT depends here on the concentration of TGF-β. These updates of the model are discussed in detail in Section 3.4.

### 3.2. The role of CAFs

An additional biological component to the invasion-metastasis cascade are the Cancer-Associated Fibroblasts (CAFs). CAFs are known to reconstruct/re-model the ECM, secrete MMPs to the extracellular environment [27, 28, 29], and are one of the main producers of TGF-β, which is responsible for the EMT procedure and will be discussed in extent in Section 3.3. We assume that the CAFs perform biased random motion, modelled via a nonlinear diffusion and haptotaxis equation, towards areas of lower ECM density. It is furthermore assumed that CAFs proliferate in a logistic manner that their proliferation is increased by the presence of ECCs, and that they die at a constant rate. Therefore, the evolution equation of the CAFs is given by

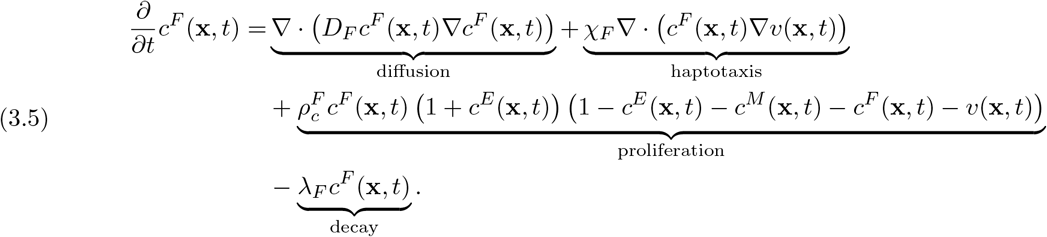

The following argument justifies the proliferation term: in the absence of ECCs, the CAFs self-proliferate in a logistic volume-filling manner, in which they compete for resources and space with themselves, ECCs, MCCs and ECM. However, we also assume that more CAFs are produced in the presence of ECCs. This accounts for the transdifferentiation of ECCs into CAFs, as well as the recruitment by ECCs of healthy fibroblast cells and their subsequent transformation into CAFs. Therefore, in areas of high ECC density, we would also expect high CAF density. Other models have explicitly included transdifferentiation [25], or healthy (non-activated) fibroblasts which then become activated to produce CAFs [30]. However, for simplicity, it is assumed that these effects are captured by the modification to the proliferation term. In the case of wound healing, fibroblast cells are directed towards areas of low ECM density as they are capable of remodelling the ECM and thus helping to repair the wounded site [31]. We have assumed that this will remain the case for CAFs and have modelled this behaviour with a haptotaxis term promoting migration in the direction of lower ECM density.

### 3.3. The role of TGF-β

The cellular (re-)programming processes EMT and MET are not random, they are rather initiated through complex interactions involving extracellular signalling molecules. Specifically, the involvement of TGF-β has been recognized as critical in triggering EMT [32, 33, 34]. In this study, we focus solely on the role of TGF-β in EMT and do not consider the impact of other signalling molecules. While this is a severe simplification of the biological reality, it allows us to investigate the specific role of TGF-β in the context of EMT and MET dynamics.

From a modelling point of view, we make the assumption that TGF-β is primarily produced by CAFs, while neglecting the contribution of ECCs and MCCs. We further posit that TGF-β diffuses freely in the environment, a condition that stands in contrast to the nonlinear diffusion assumption applied to cancer cells. This choice is substantiated by the differing diffusion time scales that molecular components—unlike cellular ones—exhibit within the tumour microenvironment. Additionally, we assume that the decay of TGF-β occurs at a constant rate.

Hence, the governing equation for the spatiotemporal evolution of TGF-β density is given by

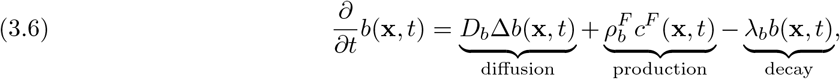

with constants 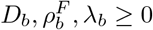.

### 3.4. The updated EMT and MET operators

Another major extension in the model concerns the EMT and MET operators in (3.4), which play a crucial role in capturing the transitions between epithelial and mesenchymal phenotypes. In the previous iteration of the model, the corresponding operators were significantly simplified. However, weit is understood that EMT and MET are complex processes influenced by a multitude of factors, including cellular heterogeneity, stochasticity, and the presence of signalling molecules. Specifically, in the current paper, we have incorporated the influence of TGF-β, which has been widely implicated in regulating EMT and MET dynamics in cancer. The updated EMT and MET operators account for the stochasticity inherent in these transitions, allowing for a more realistic representation of the phenotypic plasticity observed in cancer cells.

#### EMT

The biochemical mechanisms governing EMT remain unclear and, accordingly, significant simplifications of the modelling assumptions need to be made. Nevertheless, we assume in the current paper that EMT depends directly on the local concentration of TGF-β and, in addition, that it takes place in a random fashion. Namely, we assume that a minimum concentration of TGF-β is necessary for EMT to occur. Still, this or even higher levels of TGF-β are not sufficient to trigger EMT. To account for this uncertainty, we make the modelling assumption that EMT occurs in a random fashion.

Overall, we denote by 𝒯^EMT^ the amount of ECCs transitioning to MCCs as a time-dependent Poisson random variable

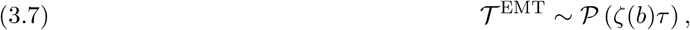

where the probability density function of 𝒫 with an arbitrary rate *λ*, is written as follows

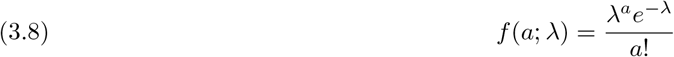

In the case of EMT, we consider a rate *ζ*(*b*) which depends directly on the concentration of TGF-β in a time interval *τ*. In other words 𝒯^EMT^ accounts for the probability that *a* events will happen in the time interval *τ*. The rate *ζ* is described as a shifted logistic function

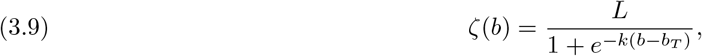

where *L, k, b*_*T*_ ≥ 0 are constants and *b* is the density of TGF-β. We choose the shifted logistic function as a switch mechanism that will ensure the necessary condition for EMT to happen, meaning that whenever *b > b*_*T*_ the rate *ζ* becomes strictly positive and reaches its highest value *L*. The values of 𝒯^EMT^ will fluctuate at different times during the evolution of the system, since the rate *ζ* takes values in the interval [0, *L*], giving us either zero when the threshold *b*_*T*_ is not exceeded, or a proportion of ECCs that undergo EMT.

We further consider that ECCs tend to undergo EMT only at times when the density of the solid tumour *c*^*E*^ is sufficiently large. Hence, we account for this restriction by multiplying 𝒯^EMT^ by a characteristic function of the form:

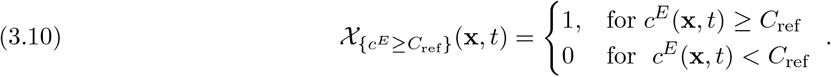

#### MET

Along with EMT, its inverse cellular programming MET, is considered to be a driver in the dissemination of carcinomas through the formation of metastases in the primary and secondary locations of the organism. The occurrence of MET depends on the type of cancer cells and tissues under investigation, and has become a target of various clinical trials, [35]. Due to the lack, though, of a clear biochemical triggering mechanism for MET, and for the sake of simplicity of presentation, we do not include in the current work any dependence on extracellular cues such as TGF-β. We will, though, assume that every solitary MCC undergoes MET randomly and independently from the others. More specifically we will assume that MET occurs in any given MCC according to the Poisson random process with a fixed rate *r >* 0 for any time interval *τ* i.e.

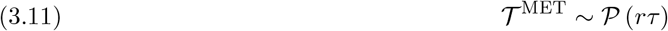

where *r >* 0 is constant.

For each one of the solitary MCCs in the subsystem (2.19), a decision is made, by (3.11), whether it will undergo MET. Then, with the use of the solitary-cell-to-density operator ℱ, the system is updated as shown in (2.21). We account for this in a similar way as we did for EMT, by multiplying 𝒯^MET^ in (3.4) by a characteristic function as in (3.10).

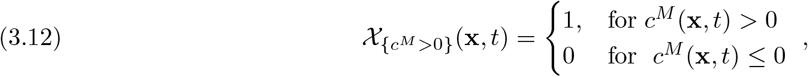

where *c*^*M*^ is the density formulation of MCCs.

### 3.5. Solitary-cell invasion SDEs

We reformulate the solitary-cell submodel so that the SDEs involve more of the biological properties describing the movement of MCCs in the tissue. The ECM plays a crucial role in the migration of solitary MMCs since MCCs move towards regions with a higher density of ECM. This implies that the velocity of MCCs is proportional to the gradient of the density of the ECM. Taking into account the haptotaxis response of the MCCs, and disregarding random effects for the moment, we write the following Ordinary Differential Equation (ODE) for the position, **X**(*t*) = **X**_*t*_, of each solitary cancer cell such as:

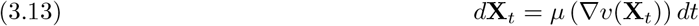

where *v* denotes the density of the ECM. The drift force now directly depends on the gradient of the ECM through the function *μ*. The form of this drift force needs to take into account that there is an upper limit in the migration speed of the cells, regardless of the density of the ECM. We denote this maximum cell speed by *V*_thr_. This function *μ* could take the following form:

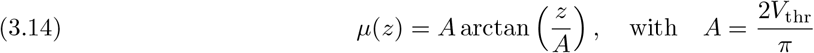

which attains values from [*−V*_thr_, *V*_thr_]. Furthermore, we reformulate the (deterministic) ODE (3.13) to an SDE in order to account for the random motion of the cells as follows:

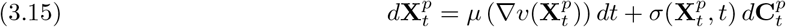

for *p* ∈ *P*, where 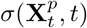 is the diffusion coefficient and 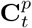 is a Compound Poisson Process (CPP). The CPP is defined as follows:

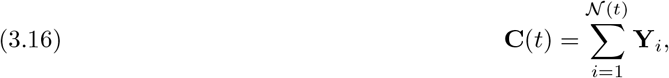

where 𝒩(*t*) is a homogeneous Poisson process with rate *λ >* 0 and 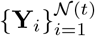 are Independent and Identically Distributed (i.i.d.) random variables. The Poisson process is a counting process that represents the number of independent discrete events that have occurred up to a time *t*. It is furthermore stationary, i.e. the probability of a random cell turning in a given time interval is the same for all equal-length intervals. Due to these properties, we find that the CPP describes the random turning in the migration of the cells better than the standard Wiener and hence employ it in (3.15).

#### Numerical solution of the SDEs

For the numerical scheme of the new SDE (3.15), we will use an Euler-Maruyama type scheme that takes the following form:

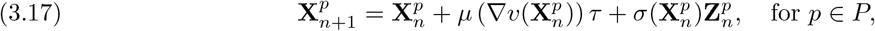

where *μ* is the drift coefficient given in (3.14) and *σ* the diffusion coefficient. For every timestep *τ >* 0 it holds

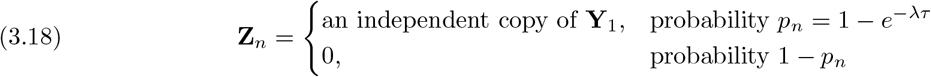

with *λ >* 0 the rate of the homogeneous Poisson process 𝒩(*t*) defined in (3.16) and **Y**_1_ a random variable. For more details on the derivation and convergence of the method (3.17) we refer the reader to [36]. We note that in every timestep of our numerical scheme, it is necessary to ensure that the cells do not exceed the maximum velocity, denoted as *V*_thr_. This situation may arise, for instance, when the combined effects of the drift and diffusion terms in (3.15) exceed *V*_thr_. In cases where ∥**V**_*t*+*τ*_ ∥ *> V* thr, with **V**_*t*+*τ*_ representing the velocity of MCC at time *t* + *τ*, we adjust the velocity to comply with the maximum limit. This adjustment is done by rescaling the newly calculated velocity as follows:

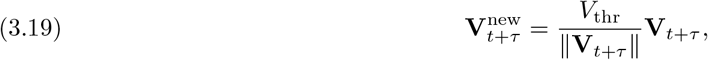

Here 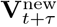 is the new, rescaled velocity that replaces the original velocity as computed by the scheme.

### 3.6. Multiple organ metastatic framework

In the final proposed extension of the mathematical model, we introduce a complex multiple-organ metastatic framework. Within this framework, an organism is conceptualized as a network of organs, interconnected by a circulatory system that serves as a conduit for cancer cells in their metastatic spread to secondary sites in the organism.

The model encompasses several critical biological phenomena, beginning with the formation of ECC-like tumour in the primary location within an organism. Following the occurrence of EMT, MCC-like cancer cells emerge in the primary organ. These MCCs migrate within the tissue and, with some probability, they infiltrate the circulatory system in a process known as intravasation. We model intravasation by a Poisson process in a similar fashion as in the EMT and MET.

Once in the bloodstream, the cancer cells acquire the designation of Circulating Tumour Cells (CTCs). The hemodynamic environment of the circulatory system imposes significant stresses upon these cells, leading to a substantial rate of destruction among the CTCs. The survival probability of the CTCs is estimated to be 0.1%, [37, 38]. In mathematical terms, we model the death rate of the CTCs through a uniform probability distribution.

CTCs that survive the circulatory network, extravasate into one of the downstream—with respect to the blood flow—organs. The anatomical arrangement and the hemodynamics of the circulatory system play a significant role in determining the sites where cancer cells may extravasate. Still, for the sake of modelling simplicity, we do not account the blood flow in this paper nor do we limit the extravasation process to organs in the downstream direction of the blood flow. We rather model the extravasation process as a Poisson event, with the choice of the extravasation location following a uniform distribution with respect to the various organs that comprise the virtual organism.

Upon arrest within the secondary location, CTCs typically undergo a phase of dormancy, [39]. For the sake of simplicity of presentation though, we do not account for cell dormancy in this paper. We instead opt to treat the newly arrested cancer cells as MCC-like, having in effect the potential to migrate in the tissue or undergo MET. This results in the formation of newly developed ECC-like densities and the engenderment of new tumours; metastasis has occurred.

In addition to the aforementioned simplifications, the geometrical representation of the various organs considered in this paper has been abstractly described as simple 3D polyhedrons of variable density. However, our multiple organ modelling framework could also account for more realistic organ geometries and structures, we refer for instance to [40, 41, 42].

## 4. The updated model

In this section, we present the fully updated model by combining all the updates introduced above in Section 3, and we also incorporate the final two components of the model—the macroscopic equations of MMPs and ECM. These system components have undergone minimal changes in the new model.

### 4.1. Density formulation

The density equations for the updated ECCs, CAFs and TGF-β have been derived in detail above and are given by (3.4), (3.5) and (3.6) respectively. To complete the updated density submodel, we must also include the density equations for the MMPs and ECM.

#### MMPs density

As before, we assume that the MMPs move through the environment via linear diffusion (molecular diffusion) and decay at a constant rate. However, with the addition of CAFs to the model, and because CAFs are known for one of the main producers of MMPs in the TME [27, 29], we now assume that CAFs are producing MMPs alongside with ECCs. Thus, the evolution equation for MMPs density is given by

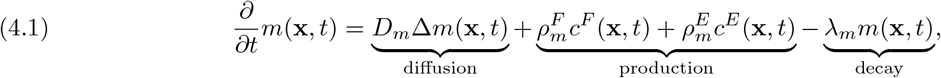

with constants *D*_*E*_, 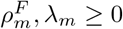. Note that equation (4.1) describes the evolution of soluble MMPs that are secreted in the local extracellular microenvironment and we do not account in (4.1) for membrane-bound MMPs such as MT1-MMP [2, 43, 44]. Although, the presence of MT1-MMP has been suggested in [43], to be a necessary and sufficient factor for the migration of MCCs. In order to separate the different types of MMPs we will only account for the membrane-bound MT1-MMP, implicitly, through the new degradation term of the ECM, shown in equation (4.2)

#### ECM density

For the new density equation of the ECM, we have added a production term with a constant rate that depends on the density of CAFs, since they are responsible for the reconstruction of the ECM [29, 45]. The degradation of the ECM is affected again by the ECC-MMP complex, instead of the MMPs alone, and we have changed the contribution of MCCs due to the effect of the membrane-bound MT1-MMPs. Hence, based on these assumptions, the evolution equation for the ECM density is given by:

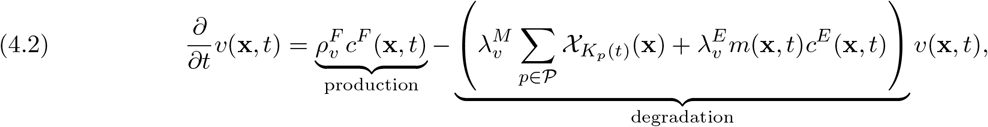

with constants 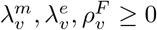. Here, *K*_*p*_(*t*) represents the physical space occupied by the mesenchymal-like cell with index *p* and 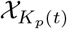 is the characteristic function defined in (2.2). The degradation of the ECM is directly impacted by the MCCs of the solitary cell subsystem rather than the density description of MCCs in (2.4).

#### Overall density submodel

Combining all the updated equations, the new overall density submodel can be written as a system of PDEs given by

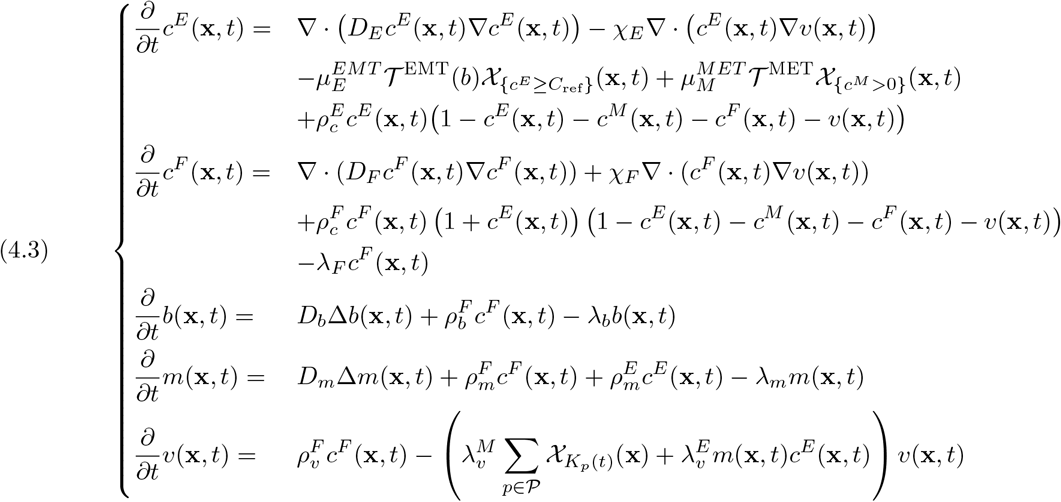

### 4.2. Solitary-cell formulation

As in the original model, the MCCs are described via a system of *N* solitary cells, indexed by *p* ∈ *P* = {1, 2, …, *N*}, where *N* can vary in time. Both the position **x**^*p*^(*t*) and the mass *m*^*p*^(*t*) are accounted for and thus the set of solitary cells is again described by (2.5). However, the evolution of MCCs is now described by the SDE (3.15), which is introduced in detail in Section 3.5.

#### Modelling reactions of MCCs

Similarly to the original model, the solitary MCCs participate in several reaction processes. These are EMT, MET, proliferation of ECCs and proliferation of CAFs. However, the new solitary-cell submodel still contains no reaction terms. We account for them in the following ways. A certain proportion of ECCs, determined by the local concentration of TGF-β, undergo EMT according to the updated EMT operator introduced in Section 3.4. This proportion of ECCs is then removed from the ECC density and added to the solitary MCC set via the density-to-solitary-cell operator which remains the same as in Section 2.3. As stated in Section 3.4, the MET operator is described by a Poisson process and so a random number of MCCs undergo MET. These solitary cells are then turned into ECC density via the solitary-cell-to-density operator given in Section 2.3. Furthermore, since the density of MCCs participates in the proliferation term for the ECCs and CAFs, at every time step the entire collection of MCCs must be transformed into a density distribution.

## 5. Numerical experiments

In this section, we conduct computer simulations using MATLAB, [46], to explore the qualitative dynamics of the updated model described in Section 4. The specifics of the numerical method used for solving the system is not discussed here. For complete details, please refer to Appendix A and B. Before initiating the simulations, we set appropriate values for all model parameters and establish suitable initial and boundary conditions. We then perform four distinct computational experiments. Through a series of plots, we visually track the system’s changes over successive time intervals. These experiments aim to demonstrate how the updated model behaves under various scenarios. However, more comprehensive studies are necessary to fully grasp the impact of the model’s new features on the process of cancer invasion.

### 5.1. Parameterization

The parameter values used in our simulations are detailed in Table 1. Whenever possible, we have sourced these values from existing literature to align our simulations with biological data. However, in several instances, it was not feasible to obtain specific values from literature, necessitating our estimation of some parameters. All values are presented in units of days, which aligns with the timescale over which our simulations are conducted.

**Table 1:** Parameter values used to perform simulations of the model.

| Symbol | Description | Value | Reference |
| --- | --- | --- | --- |
| $D_E$ | EC diffusion coeff. | $8.64 \times 10^{-8} \text{ cm}^2 \text{ d}^{-1}$ | [47] |
| $D_F$ | CAFs diffusion coeff. | $10 D_E$ | our estimate |
| $D_b$ | TGF-β diffusion coeff. | $10^3 D_E$ | our estimate |
| $D_m$ | MMPs diffusion coeff. | $10^3 D_E$ | our estimate |
| $\chi_E$ | EC haptotaxis coeff. | $10^{-3} \text{ cm}^2 \text{ d}^{-1}$ | our estimate |
| $\chi_F$ | CAFs haptotaxis coeff. | $10^{-3} \text{ cm}^2 \text{ d}^{-1}$ | our estimate |
| $\rho_c^E$ | EC proliferation rate | $1.2 \text{ d}^{-1}$ | [47] |
| $\rho_c^F$ | CAFs proliferation rate | $10^{-3} \text{ d}^{-1}$ | our estimate |
| $\rho_b^F$ | TGF-β production by CAFs | $10^{-3} \text{ g d}^{-1}$ | our estimate |
| $\rho_m^F$ | MMPs production by CAFs | $10^{-3} \text{ g d}^{-1}$ | our estimate |
| $\rho_v^F$ | ECM production by CAFs | $10^{-3} \text{ g d}^{-1}$ | our estimate |
| $\lambda_F$ | CAFs apoptosis rate | $2.62 \times 10^{-2} \text{ g d}^{-1}$ | our estimate |
| $\lambda_b$ | TGF-β decay rate | $2.62 \times 10^{-2} \text{ d}^{-1}$ | our estimate |
| $\lambda_m$ | MMPs decay rate | $2.62 \times 10^{-2} \text{ d}^{-1}$ | our estimate |
| $\lambda_v$ | ECM degradation rate | $1.8383 \text{ M cm}^{-3} \text{ d}^{-1}$ | our estimate |
| $\sigma_v$ | MC velocity diffusion coefficient | $30 \text{ cm d}^{-1/2}$ | our estimate |
| $\mu_v$ | MC velocity drift coefficient | $100 \text{ cm}^2 \text{ d}^{-1}$ | our estimate |
| $L$ | EMT maximum rate | $5 \times 10^{-3} \text{ d}^{-1}$ | [30] |
| $k$ | EMT saturation factor | $10^6 \text{ cm}^3 \text{ d}^{-1}$ | [30] |
| $\mu_M^{MET}$ | MET rate | $10^{-1} \text{ d}^{-1}$ | our estimate |
| $b_T$ | TGF-β threshold | $10^{-3} \text{ g cm}^{-3}$ | [30] |
| $s$ | maximum MC speed | $2.16 \text{ cm d}^{-1}$ | [48] |
| $v_{\max}$ | maximum ECM density | $1.06 \text{ g cm}^3$ | [49] |
| $v_{\min}$ | minimum ECM density | $0.9 v_{\max}$ | our estimate |

The computational domain, denoted as Ω, is defined as [*−*0.05, 0.05]^3^. This represents a cuboid with each side measuring 0.1cm and a total volume of 0.001cm^3^. All components of the density model are confined within this domain. Consequently, we apply homogeneous Neumann boundary conditions, which are specified below:

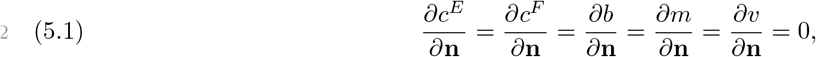

where **n** is the outward unit normal vector to the boundary of the domain Ω. In the numerical simulations shown in Section 5 we have assumed that the diffusion coefficient 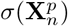 is the same for all cells and remains constant for all times. We have in addition employed a reflective boundary condition, according to which MCCs that escape the domain are returned to their last known position inside the domain.

#### Experiment 1 — Haptotaxis Flow

The first experiment we consider focuses solely on the process of EMT and the subsequent haptotaxis of the solitary MCCs. The parameter values which relate to EMT as well as MCC migration can be seen in Table 1. Further to these dynamics, we consider only the diffusion of ECCs, hence the advection and diffusion coefficients of the other model components are set to zero. Moreover, no MET takes place in this experiment, nor do any other reactions or interactions between the model components occur. The remaining initial conditions and experimental setup are described below.

First, we assume that there are no MCCs at the initial time. Second, we assume a spherical tumour of radius 0.01cm, comprised only of ECCs, is located in the centre of the domain. Within the initial tumour, we assign the density of ECCs to be 3 g cm^*−*3^ and outside this tumour the density of ECCs is set to zero. This can be written in terms of a characteristic function, see also (2.2), as

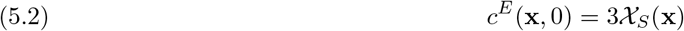

where the set *S* is given by,

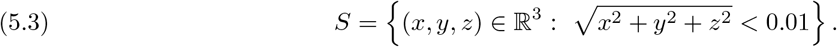

Third, CAFs are assumed to be present in a randomly selected 30% of the full domain. Over this subdomain, *Y*, the density of CAFs is uniformly distributed over the interval [0, 0.001].

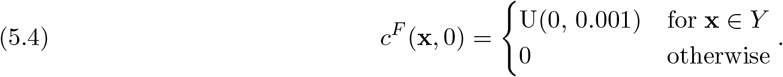

Fourth, It is assumed that initially TGF-β is present only in the area surrounding the tumour, i.e., where no ECCs are present. The density is then set according to a uniform distribution over the range [0, 0.01].

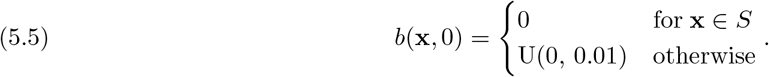

Fifth, It is assumed that MMPs are present in the whole domain, and their density is assigned a value based on a uniform distribution over the interval [0, 0.0001]

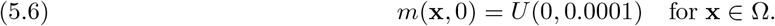

Finally, since the main aim of this experiment is to observe/verify the haptotaxis migration of MCCs, we devise an ECM that is directional with a density gradient increasing towards one of the corners of the domain. Namely, we set

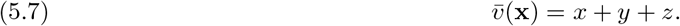

Subsequently, the ECM density values are normalised

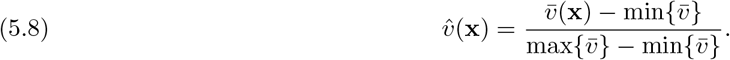

and, lastly, are brought within a biologically realistic range [*v*_min_, *v*_max_]

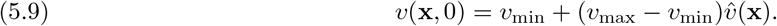

This experimental setup is largely based on an experiment carried out in [1], which we now extend to three dimensions and introduce the model updates.

From Figure 3 we observe two key properties of the model. The first is that, as EMT takes place, an initial density of ECCs is transformed into a number of individual solitary MCCs. Secondly, MCCs undergo haptotaxis, which directs their migration into areas of higher ECM density. The updated SDE, has preserved the haptotatic response of MCCs, which is encoded into the model via the advection term in the SDE (3.15), while the new random component maintains the desired qualitative behaviour of a biased random migration of MCCs. The more detailed modelling of EMT ensures that the process predominantly occurs on the periphery of the tumour. This is because the concentration of TGF-β is initially set to zero within the tumour, and EMT can only occur where the TGF-β threshold is exceeded.

**Fig. 3:**
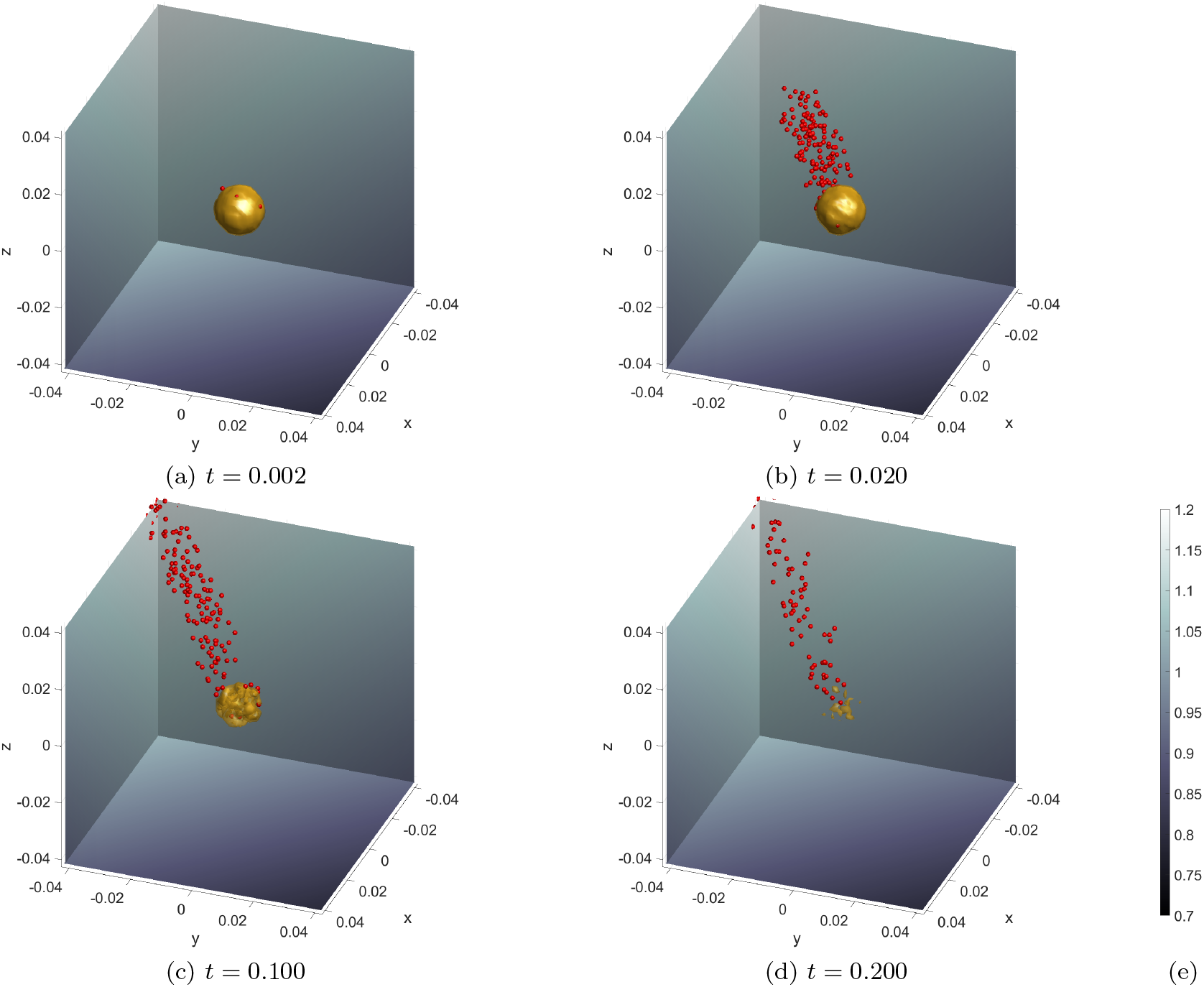
Simulation results for **Experiment 1 — Haptotaxis Flow**. Shown here is the time evolution of an initial ECC density (yellow isosurface at density value 1) depleted by EMT and the resulting solitary MCCs (red dots) migrating within the ECM (grey background). The ECM has a clear gradient towards the negative *x* and *y* and positive *z* direction (5.7)-(5.9). (a) EMT takes place in an initially spherical ECC tumour giving rise to a number of MCCs that escape from the main body of the tumour. (b) The newly formed MCCs perform a persistent random motion and migrate within the tissue towards higher ECM densities, i.e. in the direction of decreasing *x* and *y* and increasing *z*. (c)&(d) As EMT continues, more and more ECC density is depleted until the initial tumour is completely transformed into solitary MCCs. (e) Colourbar for the ECM density, common to all subplots.

#### Experiment 2 — Cancer cell islands

In this experiment, our aim is to reproduce the biologically observed phenomenon of cancer islands forming away from the main body of the primary tumour. Therefore, in this simulation, we consider the full dynamics of the model. All the parameters used in this simulation can be retrieved from Table 1. In particular, unlike the previous experiment, we now consider the process of MET, which occurs with a fixed rate given in Table 1.

The initial conditions for the ECCs, CAFs, TGF-β, and MMPs, remain the same as in Experiment 1 (5.2)-(5.6). However, the initial ECM density is set to randomly vary over the domain according to a procedure developed in [2], where we refer the reader for full details.

In the current paper we only provide a brief description of this process and refer to Figure 4 for a graphical representation. To begin with, an 8 *×* 8 matrix is created with entries taken from a standard normal distribution, *N* (0, 1). A number of refinement steps are taken until the resolution of the matrix reaches the desired (computational) resolution of the domain. At each stage, the size of the matrix is doubled to increase the resolution of the ECM. The entries to the new larger matrix are obtained from interpolating the values of the previous smaller matrix with the addition of a small amount of Gaussian noise. Therefore, as the ECM is refined it preserves the initial randomly chosen structure observed in the 8 *×* 8 matrix, with areas of higher or lower densities appearing in the same regions of the grid no matter what resolution is used. This procedure is extended into three dimensions. However, due to the increased computational time that three-dimensional simulations impose, a maximum refinement of 64 *×* 64 *×* 64 will be used for the ECM in this experiment.

**Fig. 4:**
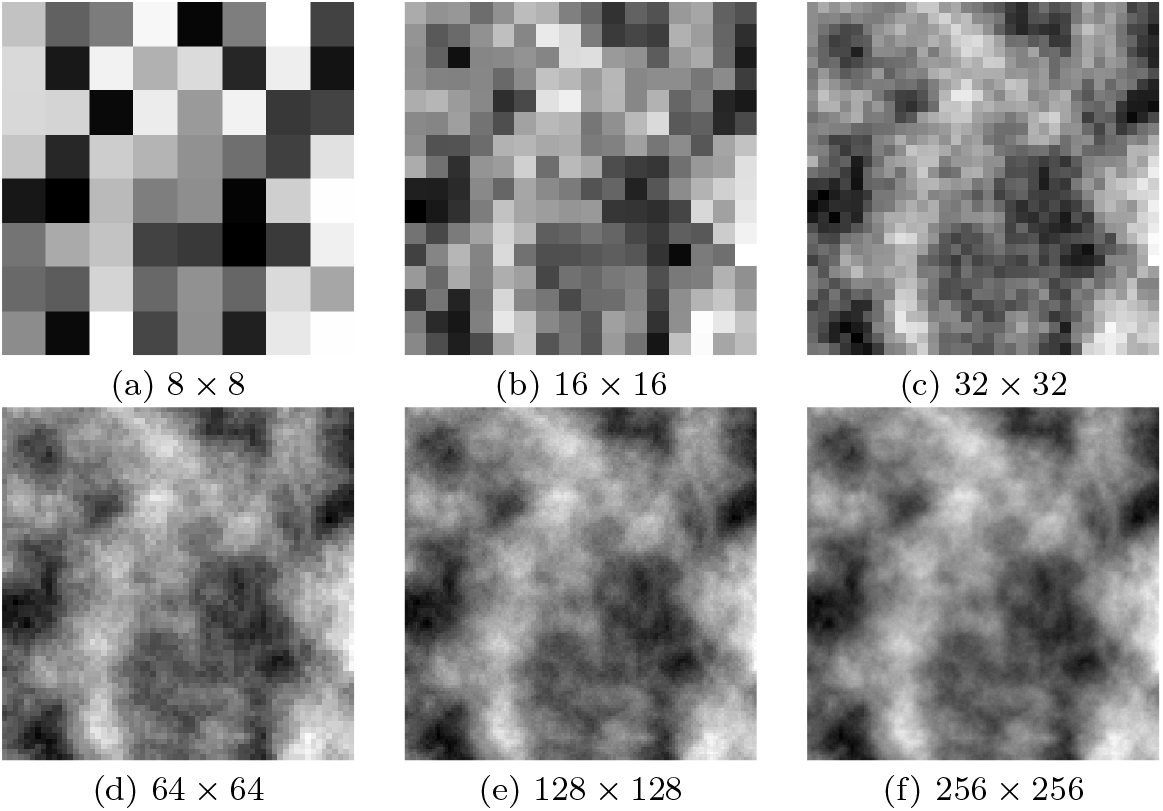
Construction of the (initial condition) ECM density employed in Experiments 2, and 4. The process starts with random values over an 8 *×* 8 grid; these introduce the basic structure of the ECM. In every stage of the construction process, the grid is bisected with the new values attained by averaging the neighbouring values of the previous coarser grid with the addition of some additive noise. The process stops when the required grid size is reached. At this stage, the values are scaled to the proper biological range. Figure source and reference [2].

The simulation results of this experiment are shown in Figure 5: The initial concentration of the ECCs is engulfed by the ECM, the CAFs, the TGF-β, and the MMPs. As time progresses, EMT takes place and solitary MCCs are materialised. These MCCs perform a biased random migration, they break free from the main body of the tumour and invade the surrounding tissue. The formation of solitary MCCs via EMT, their overall numbers, and their migration patterns are subject to the overall state of the model and the corresponding parameters. As the phenomenon progresses, the solitary MCCs undergo MET, according to the procedure previously described, acquire epithelial character, and the resulting ECCs start to proliferate. These newly formed ECCs are described in our model through their corresponding cell densities and give rise to ECC islands outside the main body of the tumour where they continue to grow.

**Fig. 5:**
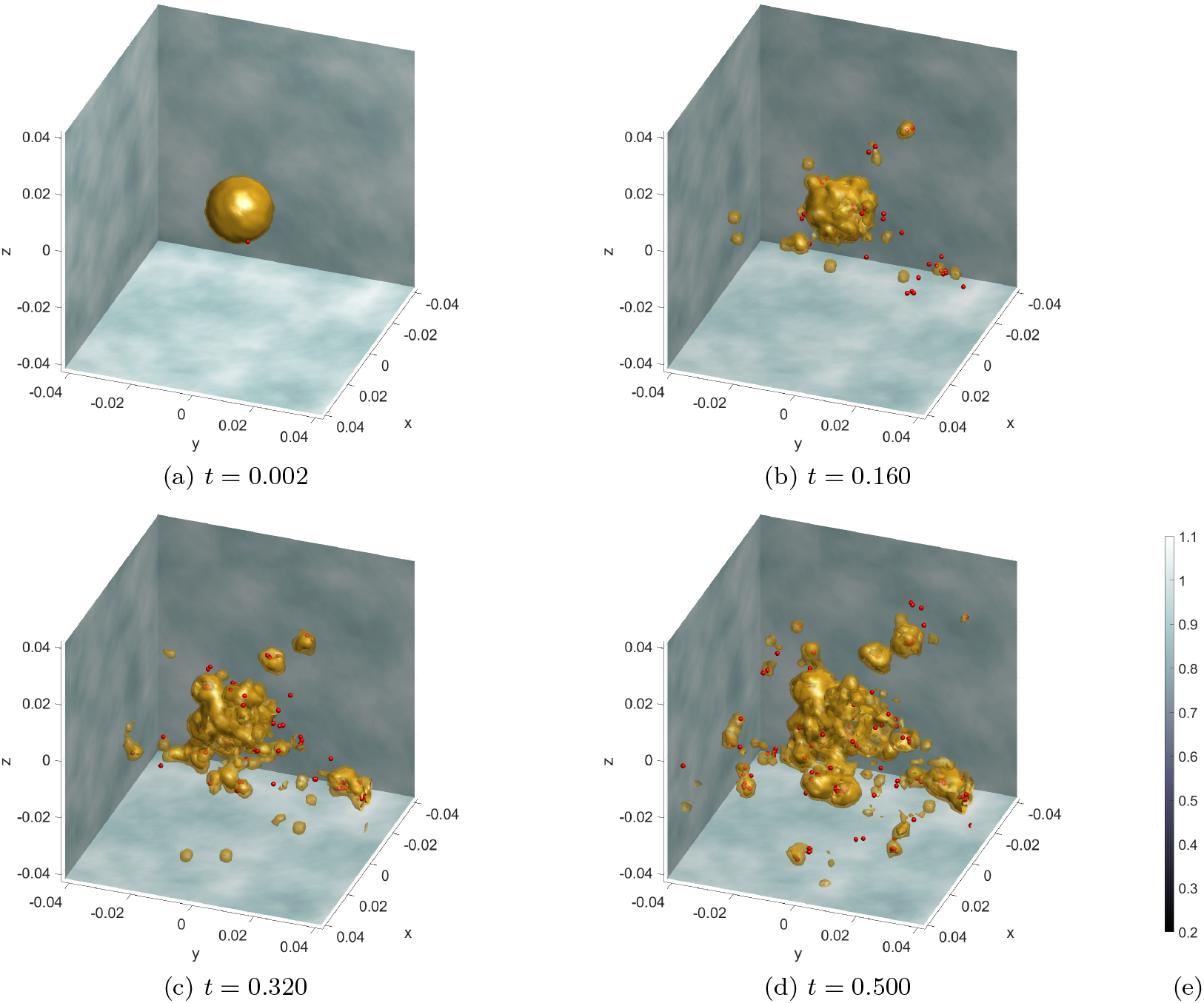
Simulation results for **Experiment 2 — Cancer Cell Islands**. Time evolution of the ECC density (yellow level 0.99 isosurface), the solitary MCCs (red points), and three slices of the ECM (grey in the background). (a) An initial spherical tumour consists solely of ECCs. (b) The ECCs undergo EMT to produce MCCs, these invade the local tissue and begin undergoing MET forming the first tumour islands. (c)-(d) The number of tumour islands increases as more MCCs undergo MET. The size of these tumour islands also increases, primarily due to the proliferation of ECCs. (e) Colourbar for the ECM density, shared by all plots.

#### Experiment 3 — Growing & Merging Microtumours

In this experiment, we investigate the complex phenomenon of (micro-)tumour merging within a specified environment. The merging process is a multifaceted event that may involve a combination of mechanical interactions, signalling pathways, and modifications in cellular behaviours. It represents a critical phase in tumour development, that potentially leads to more aggressive tumour growth and poses additional challenges for therapeutic intervention, [50].

The initial conditions for this experiment are similar to the previous ones. The main difference is that we consider two spheroids tumours rather than one as initial concentrations for the ECCs, namely,

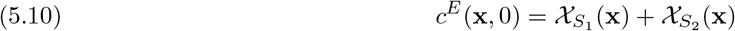

where *S*_1_ and *S*_2_ are the following translations of the set *S* given in (5.3):

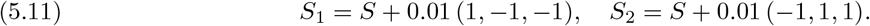

No EMT takes place in this experiment, and as no MCCs are present initially, neither does MET. Consequently, the experiment focuses solely on the behavior of ECCs, without the transformation or introduction of new cell states through these biological processes.

The simulation results are shown in Figure 6, where, as in the previous experiments, we model the primary location of the tumours as a box-like organ. Within this environment, two initially separate tumours are allowed to grow by the processes prescribed by the model (4.3). The growth of these tumours is driven by cellular proliferation.

**Fig. 6:**
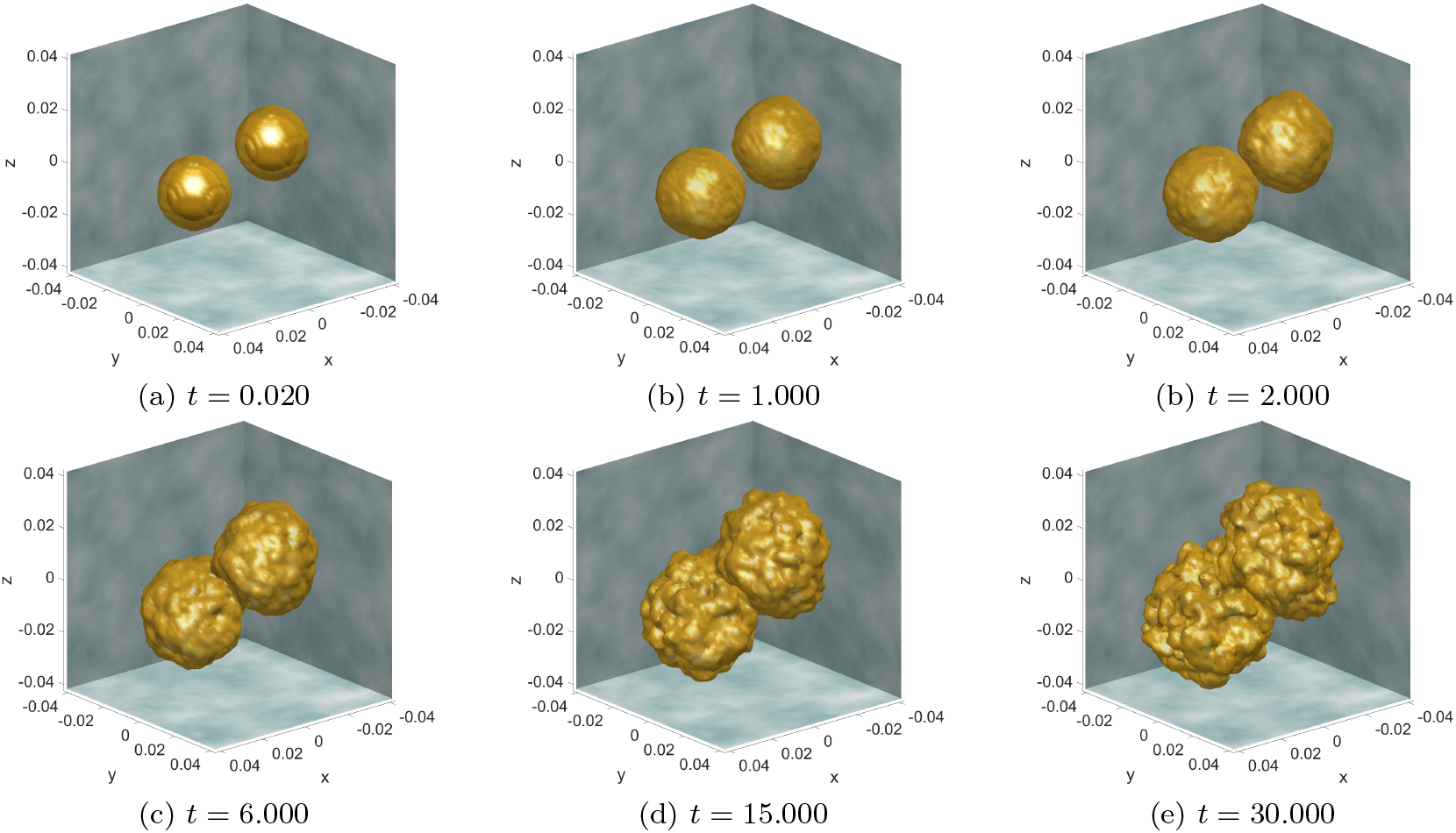
Simulation results for **Experiment 3 — Growing & Merging Microtumours**. Time evolution of two ECC densities (yellow level 0.99 isosurface) growing and merging in the absence of EMT and hence MCCs and MET.

As these tumors expand, they actively respond to the varying conditions present in the local ECM. This interaction with the ECM’s non-uniformities plays a significant role in their development and growth patterns. Over time, as these microtumors continue to grow and adapt to their immediate surroundings, they eventually reach a point where they come into contact with each other. Upon contact, the two initially separate tumors merge. This merging is a critical phase of their development, representing a new stage in the progression of the cancerous growths within the given environment. Following the merging of the two microtumors, they form a single, larger tumor. This unified tumor exhibits grows primarily along its outer surface. This is noteworthy as an indication that a tumour comprised of separate tumour islands grows faster than a single tumour of the same overall size.

This observation implies a key aspect of tumor biology—a tumor that originated from the merging of multiple smaller islands displays a more aggressive growth pattern, particularly at the periphery. This finding indicates that the internal structure and history of a tumor can have a profound impact on its growth trajectory. Understanding these dynamics is crucial for comprehending tumor development and for devising effective strategies for cancer treatment, as it provides insights into how tumors evolve and adapt over time.

#### Experiment 4 — Multiple Organ Metastasis

In this experiment, we take a further step and expand our mathematical model to account for several interconnected organs within a single virtual organism. This approach allows us to better understand how cancer progresses in the metastatic cascade.

In line with the previous experiments presented in this work, we represent the various organs as cubes of equal size. The ECM is constructed for each organ separately by the process described in Figure 4 and later employed in Experiments 2 and 3. The various organs are linked though a simplified circulatory network that facilitates the migration of cancer cells between different parts of the organism. While distinguishing between the venous, arterial, and lymphatic systems is crucial in real-world cancer progression, our model simplifies these into a single circulatory network with bidirectional connectivity, implying that cancer cells can move in any direction between interconnected organs.

In their travel through the circulatory system, the cancer cells, now termed Circulatory Tumour Cells (CTCs), are highly likely to be eradicated, with an estimated survival probability of 0.1% in our simulations.

Regarding the initial conditions and boundary conditions, the ECCs and the ECM in the organ where the tumour initially appeared are the same as in Experiment 2, with no initial ECC concentration in the remaining organs.

The simulation results, presented in Figure 7 clear exhibit the metastatic cascade as is perceived by the this modelling framework. Namely, a tumour grows in a primary location of the organism. EMT takes place leading tot he invasion of MCCs in the local tissue (first row). Gradually the MCCs enter the blood stream and spread to secondary locations in the organism. Both in the primary and secondary sites, the MCCs undergo (potentially) MET giving rise to ECCs (second row). The new formed ECC islands grow due to proliferation and give rise to a number of micrometastases (third row). These micrometastases serve as precursors to larger and and more dangerous metastases which with time merge and form larger tumours (forth row).

**Fig. 7:**
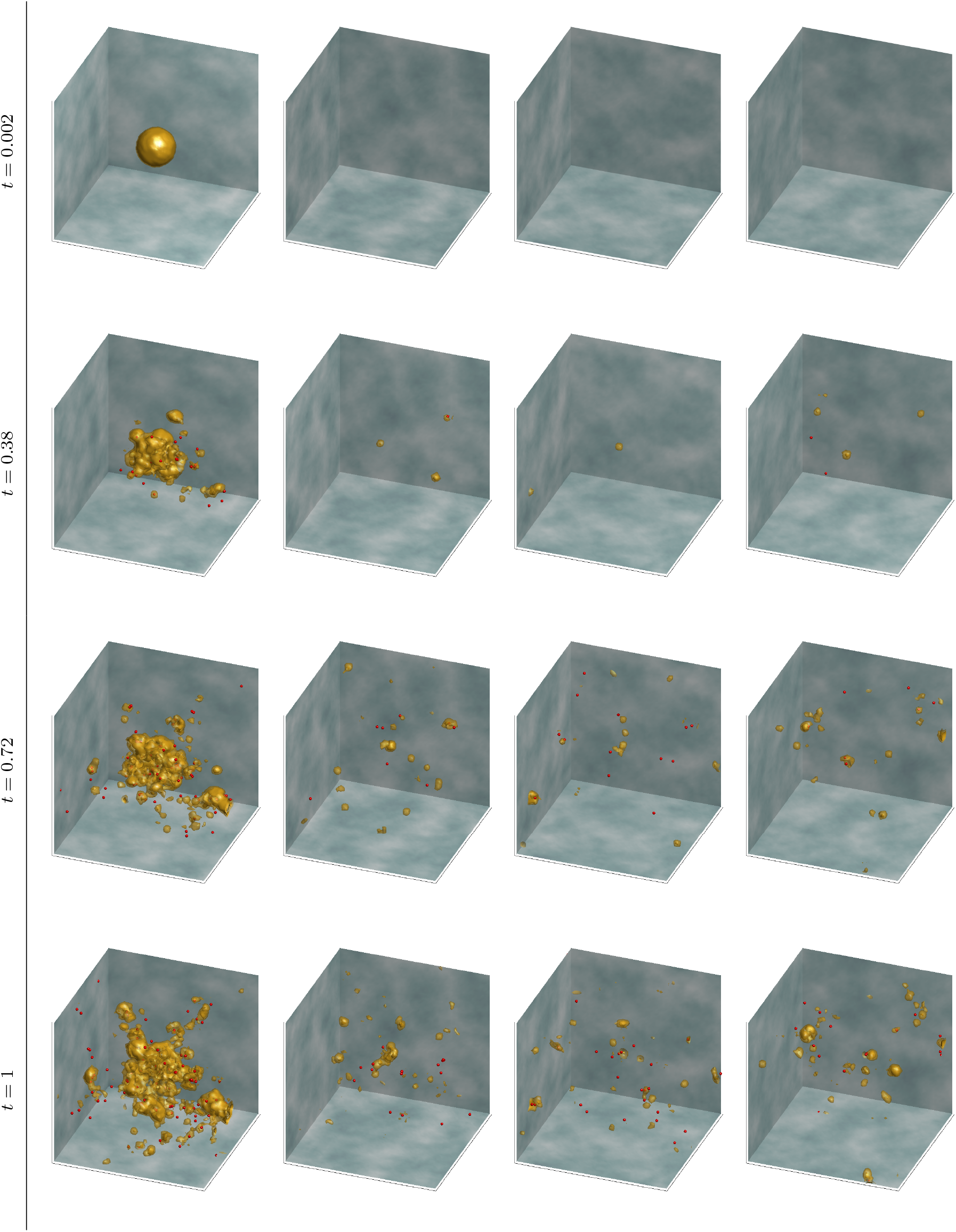
Simulation results for **Experiment 4 — Multiple Organ Metastasis**. this figure illustrates the progression (across rows) of metastatic growth in four different organs (columns). The various organs are represented cubes whose size and colour schemes are the same as in Figure 5. At *t* = 0.002: An initial ECC tumour has already developed to a notable size in the first organ (column 1), at which point EMT leads to the formation of MCCs. At *t* = 0.38: These MCCs migrate within the first organ and have beagun spreading, via the circulatory network, to secondary organs (columns 2 to 4). At *t* = 0.72: micro-metastases are observed forming in the secondary organs concurently with the growth of the primary tumour. At *t* = 1: there is a continued expansion of both the initial tumor and the developing micro-metastases, indicating progressive metastatic spread.

As noted in the previous experiment, the simulations presented here reveal again that when cancer cells initially form separate, disjoint islands, their overall volume tends to increase more quickly compared to a scenario where they start as a single, unified mass. This difference in growth rate is not just marginal, indicating a fundamental aspect of tumor biology. Understanding these dynamics is crucial for developing effective therapeutic strategies, as it highlights the importance of considering not just the size but also the distribution and structure of cancerous growths.

## 6. Discussion

In this paper, we introduce an updated version of our previously proposed genuinely hybrid multiscale tissue cancer invasion model [1, 2]. This enhanced model replicates more naturally the transition from the epithelial invasion strategy original of the ECCs to the individual invasion strategy of the MCCs. The model is comprised of a system of PDEs and SDEs that describe the progression of the ECCs and MCCs, respectively, while incorporating the phenotypic transitions of EMT and MET.

A significant upgrade in our model is the integration of nonlinear degenerate diffusion—similar to a porous medium—in the evolution equation of the ECCs and the other living cell components. This model enhancement contributes to a more realistic representation of the tumour growth in the tissue microenvironment, addressing the issue of infinite propagation speed that is commonly encountered in linear diffusion models.

Another significant extension is the updated modelling of EMT and MET. We have considered the role of TGF-β as an EMT trigger and detailed the modelling process including the inherent randomness associated with MET. Moreover, we have introduced CAF cells into our model to reflect their role in shaping the tumour microenvironment through the reconstruction of the ECM.

We have furthermore, updated the structure of the SDEs that drive the migration of MCCs. Namely, we have accounted for a Compound Poisson Process as the core modelling framework of the stochasticity in the change of direction in the migration of the cells. This approach is in contrast to the typically employed Wiener processes in problems of a similar nature and we have opted for the Compound Poisson Process for its greater biological relevance.

Another substantial extension of the hybrid model introduced in this paper is a multiple-organ metastatic framework. Namely, we have developed a basic circulatory network that links various organs, each being modelled by a “local” version of the hybrid model. This modelling framework allows MCCs to enter the bloodstream and, if they survive the stresses of the circulatory network, be transferred to secondary locations in the organism. Once there, the MCCs may undergo MET, start to proliferate, and give rise to metastatic tumours.

We have validated these model extensions through several numerical experiments presented in Section 5. These simulations demonstrate the updated model’s capabilities to reproduce phenomena that are biologically relevant and generate predictions consistent with experimental observations. They highlight the model’s potential utility in exploring specific facets of cancer biology and treatment response.

We envision further enhancements, particularly the integration of immune system interactions within the tumor microenvironment and the refinement of the circulatory system for enhanced physiological accuracy, hold great promise for advancing theoretical research and potentially clinical applications in cancer treatment. These developments will not just be incremental improvements but significant steps towards transforming our model into a comprehensive tool that can navigate the complexities of cancer biology. By more accurately simulating the interplay between tumors and the body’s defenses, as well as the spread of cancer through vascular networks, it is expected that the model will impact clinical decision-making and optimized treatment strategies. These developments represent a promising direction in cancer research, potentially leading to more nuanced insights and improved therapeutic strategies.

## Appendix A Time evolution of the two coupled cancer cell types

We provide here a brief account of the numerical method used to simulate the time evolution of the overall model. For more details, the reader is referred to [1, 2]. We denote the density and solitary-cell variables as **w**(**x**, *t*) and 𝒫^*β*^(*t*) respectively. The numerical approximation of the complete system at the instantaneous time *t* = *t*^*n*^ is therefore given by

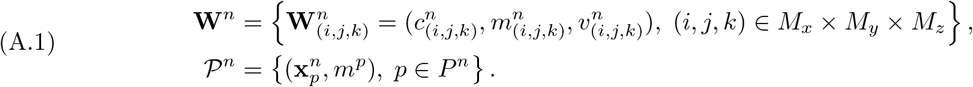

Here we extend the two-dimensional notation used in [1] to include a third dimension, where *M*_*x*_, *M*_*y*_, *M*_*z*_ ∈ ℕ denote the resolution of the numerical grid in the x-, y-and z-directions respectively.

The evolution of the density and cell variables comprises two processes: time evolution and phase transitions (EMT and MET). In particular, a *operator splitting* approach is used. That is the time period *t* ∈ [*t*^*n*^, *t*^*n*+1^], where *t*^*n*+1^ = *t*^*n*^ + *τ*^*n*^, is split into two half time steps, with phase transitions (EMT and MET) occurring in the middle. The three steps of this operator-splitting approach are outlined below.

- During the first half-time step, 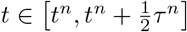, the system evolves without the influence of EMT or MET. Using tuple notation this can be written as

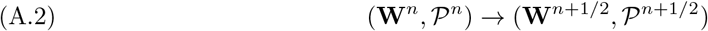

with

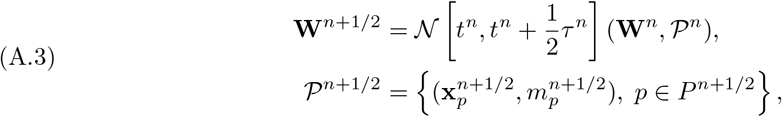

where 𝒩[*t, t* + *τ*] is the numerical solution operator responsible for the spatiotemporal evolution of the density variables. This uses an *implicit-explicit Runge-Kutta finite volume* method (IMEX-RK), which is discussed in more detail in Appendix B. The individual cells evolve according to the *Euler-Maruyama* type scheme, which is discussed in section 3.5, rewriting equation (3.17) for the half-time step gives

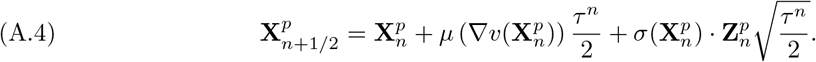

Since it is assumed that no EMT or MET takes place during this half-time step the number of solitary cells, their indices, and their masses remain the same, hence 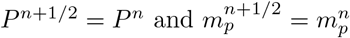.The combined spatiotemporal evolution of the density and solitary-cell variables for this half-time step can be written compactly in operator notation as

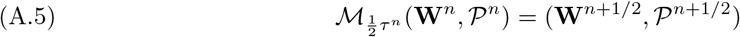
- At the midpoint of the time interval, 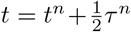, EMT and MET takes place. The phase transitions are assumed to be instantaneous and are represented by the operators ℛ^*EMT*^ and ℛ^*MET*^, introduced in (2.18) and (2.21) respectively. Combining both these processes into the single operator 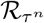, the overall system evolves as

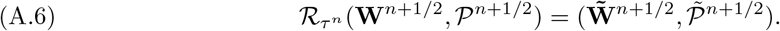
- During the second half-time step, 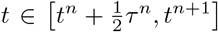, the density and solitary cells again evolve without EMT or MET taking place. This occurs in an essentially identical fashion as in the first half-time step. Using the same operator notation as for the first half-time step, the tuple of the overall system evolves as

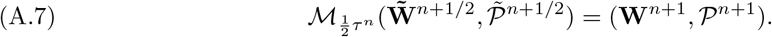

Overall combining (A.5), (A.6), and (A.7), the evolution operator for the system over a single time step can be written as a splitting method of the form

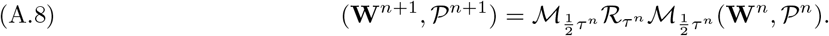

## Appendix B Numerical solution of PDE systems

The advection-reaction-diffusion system (4.3) is solved numerically using a specific second-order Implicit-Explicit Runge-Kutta Finite Differences, Finite Volumes (IMEX-RK FD-FV) numerical method. This method constitutes an extension of a previous method developed and employed in [51, 25, 40, 52, 53, 1, 2] where we refer for most of the details. Here, we only discuss some of its components.

We consider, at first, a generic advection-reaction diffusion system of the form

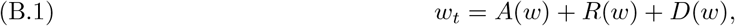

where *w* represents the analytical solution vector of the system, and *A, R*, and *D* are the advection, reaction and diffusion operators respectively. After spatial discretisations have taken place, we denote the corresponding semi-discrete approximation by *w*_*h*_, where the index *h* denotes the (maximal, if the space discretisation is non-uniform) spatial grid diameter. The semi-discrete solution *w*_*h*_ satisfies the following system of ODEs

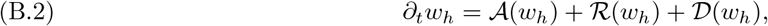

where the numerical operators 𝒜, ℛ and 𝒟 are (spatially) discrete approximations of the advection-reaction, diffusion operators, *A, R* and *D* in equation (B.1). Moreover, as the numerical scheme we employ is (partially) FV, raising its accuracy to the second order necessitates the use of flux limiters for the interface reconstruction of the numerical fluxes. Out of a large number of flux limiter options, we have found that the Minimized-Central (MC), see [54], constitutes a robust and efficient choice.

Before solving (B.2), we re-organise its terms in implicit and explicit (IMEX splitting) and, accordingly, (B.2) takes the form

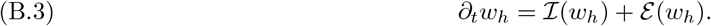

The actual IMEX splitting depends on the problem at hand, but in a typical case the advection terms 𝒜 are treated explicitly in time, the diffusion terms 𝒟 implicitly, and the reaction ℛ terms either explicitly or implicitly, depending on their stiffness. In the problems that we encounter in this paper, all reaction terms have been resolved explicitly in time.

The semi-discrete problem (B.3) is now solved using a diagonally implicit RK method for the implicit part ℐ(*w*_*h*_), and an explicit RK for the explicit part *ε*(*w*_*h*_). Altogether, we solve (B.3) using the additive RK scheme

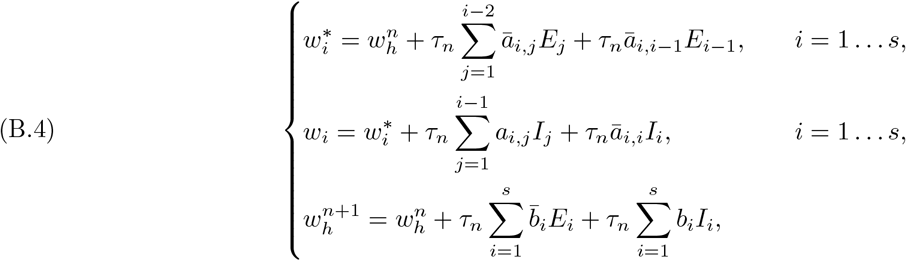

where *s* = 4 are the stages of the IMEX-RK method, *E*_*i*_ = ε(*w*_*i*_), *I*_*i*_ = ℐ(*w*_*i*_), *i* = 1 … *s*, and 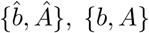 are the coefficients for the explicit and implicit part of the scheme respectively. These coefficients can be found in the Butcher Tableau in Table 2, cf. [55]. As a final stage of this method, we solve the linear system in (B.4) using the Iterative Biconjugate Gradient Stabilised Krylov subspace method, see [56, 57].

**Table 2:**
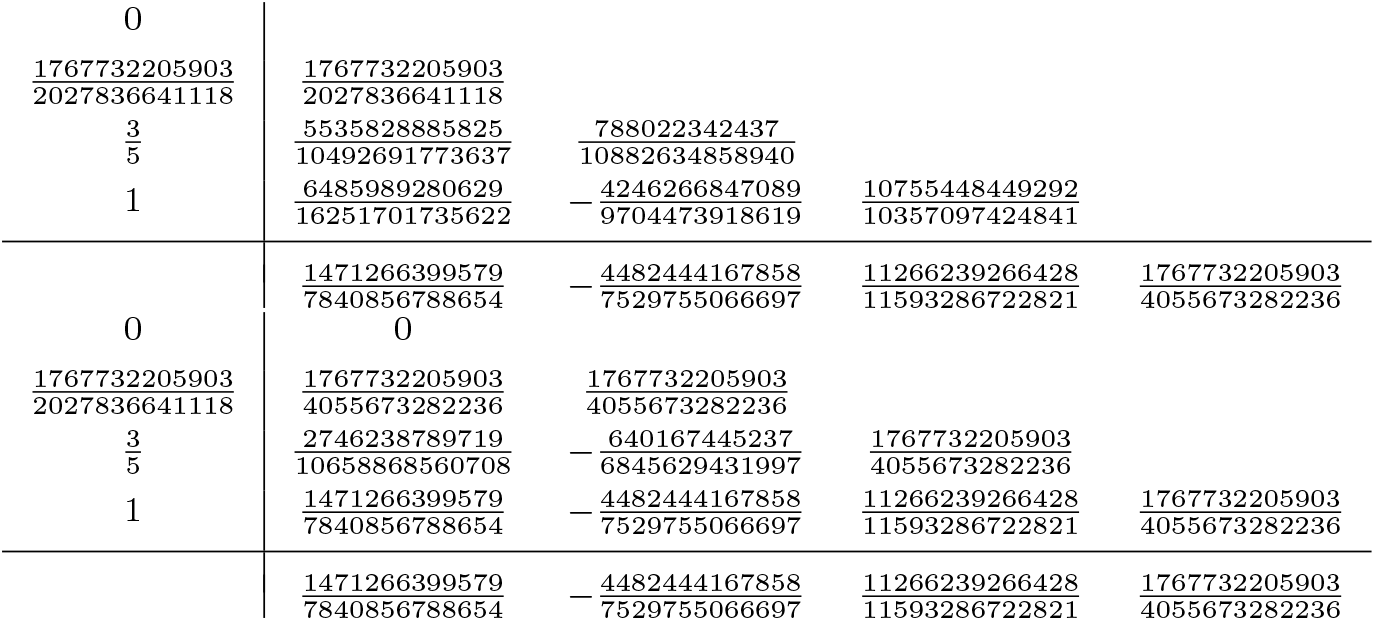
Butcher tableaux for the explicit (upper) and the implicit (lower) parts of the third order IMEX scheme (B.4), see also [55].

